# Rapid reversible RNA isoform switching during loss and recovery of turgor in *Arabidopsis thaliana*

**DOI:** 10.64898/2026.03.12.711442

**Authors:** Jazmine L. Humphreys, Luke A. Yates, Jakob B. Butler, Steven M. Smith

**Affiliations:** ARC Centre of Excellence for Plant Success in Nature and Agriculture, School of Natural Sciences, University of Tasmania, Hobart, Tasmania, Australia

**Keywords:** Arabidopsis thaliana, RNA isoform transcriptome, RNA isoform switching, alternative RNA splicing, turgor, water potential

## Abstract

Plant genes can each produce multiple RNA isoforms, but little is known about when and how they are produced, or their functions. We conducted a detailed time course analysis of rapid changes in abundance of RNA isoforms in response to transient water deficit in *Arabidopsis thaliana* seedlings. We identified 95,104 transcripts in total, 21,935 of which were differentially expressed. Strikingly, 1,258 differentially expressed genes were identified only by means of RNA isoform analysis and would have been missed by conventional analysis. Several hundred genes exhibited a very rapid and reversible switch in the most abundant RNA isoform, with timings corresponding to defined changes in seedling water potential. In most cases it is predicted that RNA isoform switching would generate new protein products. Many of these genes encoded proteins of RNA metabolism and splicing, while others potentially function more directly in the response to water deficit. We propose a model in which very rapid RNA processing mechanisms come into play within minutes of imposition of the stress, changing the RNA isoform population including RNAs that encode components of the RNA processing apparatus itself. These changes together with subsequent changes in transcription facilitate further changes in the RNA isoform population as part of the stress and recovery responses. More broadly we propose that responses to abiotic stress involve substantial changes in gene function through the production of RNA isoforms, and we present a detailed approach to identifying such genome-wide RNA isoform switching.

## Introduction

Multiple RNA isoforms can be produced from a single gene through numerous mechanisms including alternative transcription start sites, RNA splicing, editing and alternative polyadenylation sites, substantially expanding the functional repertoire of eukaryotic genomes. In *Arabidopsis thaliana*, recently reported transcriptome assemblies indicate that over half of protein-coding genes produce multiple RNA isoforms, with one reference transcriptome (AtRTD3) now annotating nearly 170,000 transcripts (Zhang et al., 2022). Many of these RNA isoforms encode proteins with distinct domain architectures, subcellular localisations, or other structural features (protein isoforms), while other RNA isoforms can potentially be targeted for transport, sequestration or destruction, such as by nonsense-mediated decay (NMD) (Tognacca et al., 2023).

The production and function of RNA isoforms has been implicated in many aspects of plant biology, including developmental programs (Staiger and Brown, 2013), responses of plants to biotic interactions (Godinho et al., 2025), and responses to abiotic stress, including drought, temperature extremes, and high light (Shikata et al., 2014; Capovilla et al., 2015; Laloum et al., 2018; Alhabsi et al., 2025). For example, *FLOWERING LOCUS M* undergoes RNA isoform switching at high temperature to produce an isoform that promotes flowering (Posé et al., 2013). The *RESISTANCE TO PSEUDOMONAS SYRINGAE 4* gene requires coordinated expression of both full-length and truncated RNA isoforms for disease resistance (Zhang and Gassmann, 2003). Under osmotic stress, alternative splicing of *PYRROLINE-5-CARBOXYLATE SYNTHASE 1* (*P5CS1)* transcripts contributes to differences in proline accumulation and drought tolerance (Kesari et al., 2012). The mechanosensitive calcium channel OSCA1.1 is implicated in osmosensing in response to turgor stress, during which an RNA isoform containing an inhibitory upstream open reading frame (uORF) declines in amount, potentially increasing OSCA1.1 production from an RNA isoform lacking the uORF (Yuan et al., 2014; Wu et al., 2024). While such studies have established that RNA isoform production is a significant component of stress responses, in the majority of cases the timing and scale of RNA isoform production, the mechanisms involved, and the functions of such RNA isoforms are poorly understood.

Climate change is expected to have an increasingly negative impact on plant growth, particularly due to increasing temperatures and variable water availability. Although alternative RNA splicing is recognised as an important component of drought responses over hours to days (Laloum et al., 2018), plants also routinely experience much more rapid, transient changes in water status. Vapour pressure deficit (VPD), determined largely by air temperature and humidity (Grossiord et al., 2020), can increase abruptly, reducing leaf water potential and turgor within minutes (McAdam and Brodribb, 2015; McAdam et al., 2016; Sussmilch et al., 2017). Stomatal regulation can limit foliar water loss, but under high VPD this compensation may be insufficient, leading to transient declines in turgor. While there is evidence that RNA isoforms play a role in longer-term drought responses (Laloum et al., 2018; Alhabsi et al., 2025), a role for RNA isoforms in rapid or transient responses to water deficit has not been reported.

The aim of this study was to identify rapid changes in RNA isoforms in *A. thaliana* seedlings that occur as a result of a rapid drop in water potential and turgor. Isolating turgor-specific responses is experimentally challenging because plants are highly sensitive to mechanical stimulation. Rapid changes in plant transcriptomes are observed within minutes in response to numerous mechanical cues including brief touching (Darwish et al., 2022; Matsumura et al., 2022), gentle shaking for a few seconds (Kimbrough et al., 2004), and even raindrops (Matsumura et al., 2022). We addressed this challenge by rapidly increasing VPD to induce turgor loss without any other physical disturbance, following which recovery of turgor occurred by water uptake through the roots. The onset of changes in transcript abundance and the subsequent trajectory of these transcripts was determined using Change Point Additive Models (*cpam*), newly developed to analyse time series data (Yates et al., 2026). The resulting dataset provides a resource for identifying turgor-responsive genes. Furthermore, the methods presented here provide a general approach to discovering genome-wide RNA isoform switching in response to abiotic stresses.

## Results and discussion

### An experimental system to study responses to changes in turgor

To characterise changes in the transcriptome in response to the rapid loss of water potential and turgor, and their subsequent recovery, we developed an experimental system in which *A. thaliana* seedlings were grown on agar nutrient medium in petri dishes for 12 d with a relative humidity (RH) of approximately 98%. By carefully removing the lids the seedlings were then exposed to a RH of 50% and thus experienced a rapid increase in vapour pressure deficit (VPD) from 0.1 to 1.2 kPa (Figure 1A). This treatment resulted in a decline in seedling water potential from −0.57 MPa to −1.9 MPa in the following 30 min, which we refer to as Phase I (Figure 1B). In the subsequent 30 min (Phase II) water potential recovered to −1.1 MPa and thereafter it stabilised at −1.1 MPa up to 240 min (Phase III) due to water uptake from the medium through the roots.

**Figure 1.**
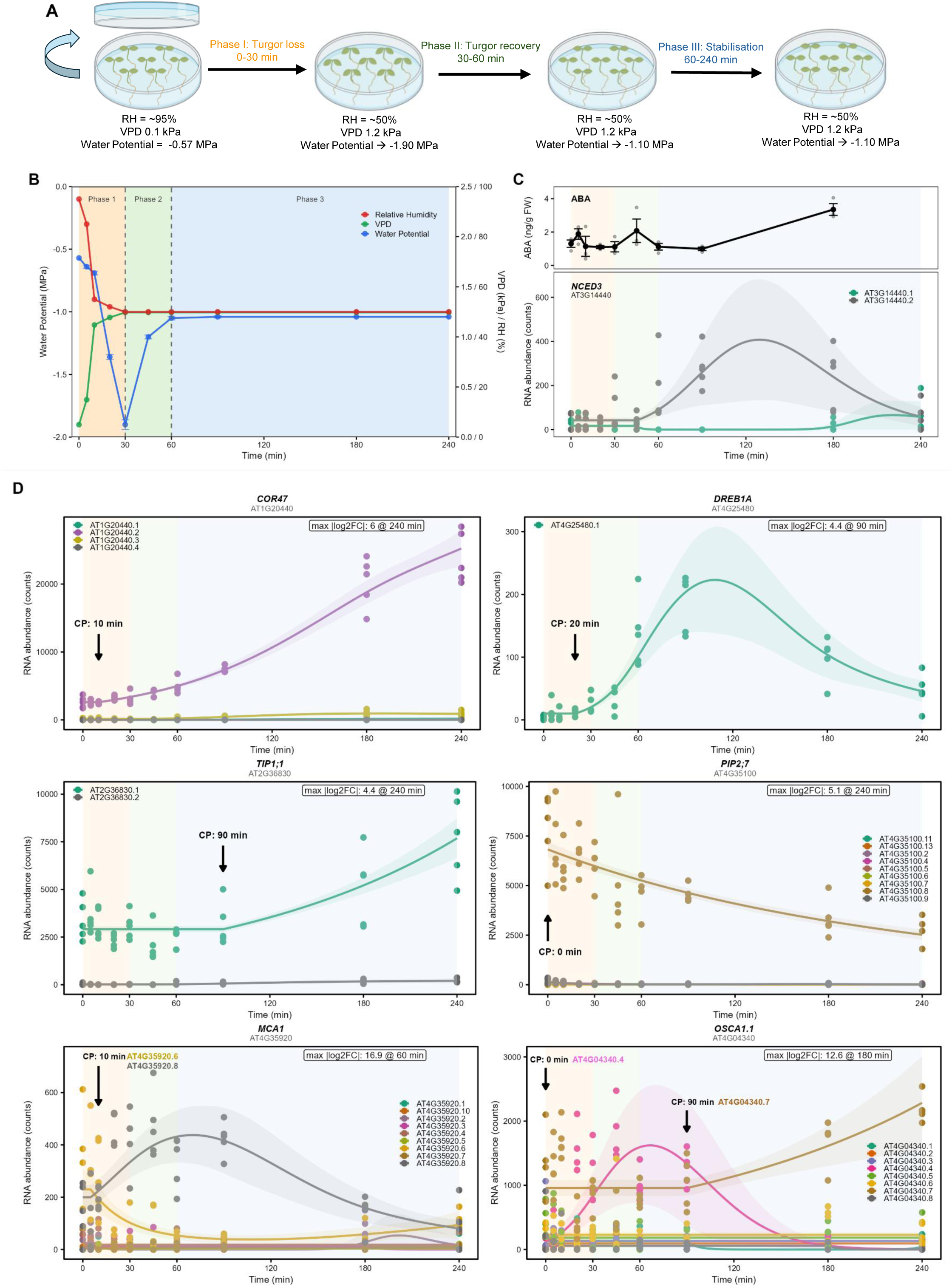
An experimental system for turgor loss and recovery. (A) Schematic of the experimental system showing changes in physiological parameters. RH: relative humidity, VPD: vapour pressure deficit. (B) Time course of seedling Water Potential (blue), Relative Humidity (red), and VPD (green). Coloured bands indicate Phase I (0–30 min; yellow), Phase II (30–60 min; green), and Phase III (60–240 min; blue). Phase boundaries are indicated by dashed lines. Data are means ± SE (n = 3 per timepoint). (C) Upper panel: abscisic acid (ABA) content (ng/g FW). Data are means ± SE (n = 3). Lower panel: *cpam*-fitted trajectories for both annotated transcripts of *NCED3* (AT3G14440.1 and AT3G14440.2). Individual data points show replicate values (n = 5 per timepoint), solid line represents *cpam*-fitted trajectory, shaded ribbons encompassing the trajectory represent 95% confidence intervals. (D) *cpam*-fitted trajectories for transcripts of six well-characterised water deficit-responsive genes illustrating distinct response patterns. Individual data points show replicate values (n = 5 per timepoint), solid line represents *cpam*-fitted trajectory, shaded ribbons encompassing the trajectory represent 95% confidence intervals. Changepoints (CP) and maximum log₂ fold-change values are indicated for each transcript. Transcript identifiers (e.g. AT1G20440.1, AT1G20440.2) follow AtRTD3 annotations, where the numerical suffix distinguishes distinct RNA isoforms from the same gene.

RNA was extracted from seedlings for sequencing at each of ten timepoints spanning all three phases, with five biological replicates at each timepoint. For each gene we determined the relative abundance of all transcripts as defined by the AtRTD3 reference transcriptome which identifies 169,503 transcripts from 40,932 genes (Zhang et al., 2022). We employed Change Point Additive Models (*cpam*) which identifies the first timepoint at which the amount of each transcript deviates significantly from baseline (the changepoint) and models the subsequent trajectory of each transcript (Yates et al., 2026). We refer to any transcript that changes significantly in abundance during the experiment as a Differentially Expressed Transcript (DET).

### Responses of genes known to be responsive to water deficit

To illustrate the key features of *cpam* and to validate the experimental system we examined transcripts of some genes known to be responsive to water deficit, including ABA biosynthesis, transcriptional regulation, water transport and mechanosensing (see Supplemental Table 1 for full names and functions). The *NCED3* gene encoding a key step in ABA biosynthesis typically increases in expression during drought stress (Sussmilch et al., 2017). This gene has one dominant transcript, which begins to increase after 45 min (the changepoint) and peaks between 90 and 120 min, after water potential has recovered (Figure 1C). A less abundant transcript exhibits the reciprocal pattern (Figure 1C). Extracts of seedlings contained very low levels of abscisic acid (ABA) with only a very small increase after 180 min (Figure 1C). These results imply that ABA does not play a major role in the rapid response of seedlings to changes in VPD and water potential.

The drought-responsive genes *COR47* and *DREB1A* (Supplemental Table 1) both initiated responses in Phase I but had distinct trajectories thereafter (Figure 1D). *COR47* has four RNA isoforms but only one DET, which showed sustained accumulation throughout the experiment. In contrast, *DREB1A* has only one transcript which displayed a transient peak in Phase III. This distinction between sustained and transient responses illustrates the value of such detailed time course analysis and suggests that COR47 and DREB1A could have different functions in the response to loss and recovery of water potential.

The aquaporin genes *TIP1;1* and *PIP2;7* produced transcripts showing opposing trends on different timescales. *TIP1;1* produced one DET which increased only after 90 min while *PIP2;7* produced eight transcripts one of which was a DET, declining progressively from the beginning (changepoint 0 min). These contrasting dynamics suggest distinct roles during initiation of turgor loss and much later stabilisation (Figure 1D).

The genes encoding mechanosensitive channels *MCA1* and *OSCA1.1* illustrate additional complexity at the level of individual genes (Figure 1D). *MCA1* produced nine RNA isoforms, two of which were DETs showing identical changepoints (10 min) but opposite trajectories, one increasing and one decreasing. The *OSCA1.1* gene produced eight RNA isoforms, two of which were DETs and showed very different changepoints and trajectories: one responding immediately (changepoint 0 min) with a transient peak, and the other initiating much later (90 min) with sustained accumulation (Figure 1D).

These results validate the experimental system by revealing rapid changes in transcript abundance from well-characterised genes known to respond to water deficit. They further reveal that for single genes, different transcripts can be produced at different times with distinct trajectories, suggesting that individual genes may contribute to the stress response in multiple ways.

### Origin and functions of DETs

The reference transcriptome (AtRTD3) annotates 169,503 transcripts from 40,932 genes. We identified 154,383 of these transcripts but only 95,104 were of sufficient abundance to justify further analysis, and of these, 21,935 were DETs (Figure 2A, Supplemental Table 2), arising from 10,167 genes. Strikingly, *cpam* analysis showed that changepoints for these DETs clustered strongly within Phase I, with the majority of transcripts initiating their response within 10 min of the change in VPD (Figure 2B). Fewer DETs had changepoints during Phase II, while an increase occurred in Phase III. This pattern indicates that most responses are triggered very rapidly during turgor loss. Of the genes annotated in AtRTD3, approximately half (20,394) produce multiple RNA isoforms (Figure 2C). Such genes were over-represented among those responding to a change in water potential (Figure 2C). Furthermore, of the 10,167 genes producing DETs, the majority (5,297) produced two or more DETs (Figure 2C), suggesting that RNA isoform complexity may be a feature of stress-responsive loci.

**Figure 2.**
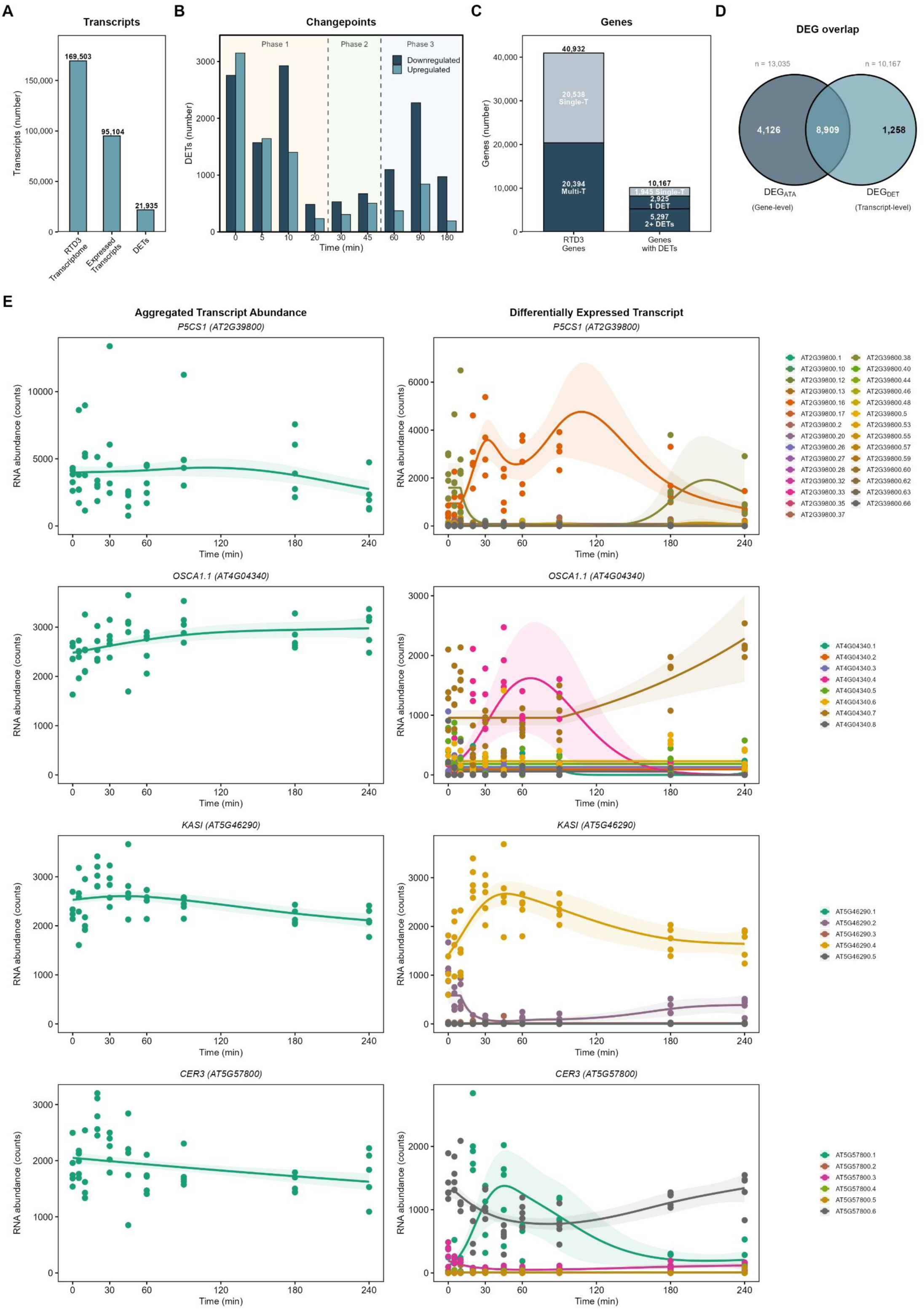
Multi-transcript genes are the major source of DETs. (A) Number of transcripts at each stage of the analysis pipeline. Annotated in the AtRTD3 reference transcriptome, Expressed Transcripts are those detected in the present study, DETs are classified as Differentially Expressed Transcripts. (B) Distribution of changepoints for all DETs across the experimental time course, separated by direction (Downregulated, dark blue and Upregulated, light blue). Phase boundaries are indicated by dashed lines. A changepoint at a given timepoint represents the onset of significant deviation from baseline between that timepoint and the next sampling point e.g. a changepoint at time 0 represents an observed change between 0 and 5 min. (C) Left: number of genes in AtRTD3 producing single (Single-T, upper, dark blue) or multiple (Multi-T, lower, light blue) transcripts. Right: Number of DETs produced from single-T and multi-T genes (right). (D) Overlap between differentially expressed genes identified by aggregated transcript counts (DEG_ATA; n = 13,035) and RNA isoform analysis (DEG_DET; n = 10,167). (E) Aggregated counts (left) and individual RNA isoform counts (right) for four example genes: *P5CS1* (AT2G39800), *OSCA1.1* (AT4G04340), *KASI* (AT5G46290), and *CER3* (AT5G57800). *cpam*-fitted trajectories are shown as described in Figure 1.

Gene Ontology enrichment analysis of DETs defined by phase and direction revealed distinct functional categories associated with each phase of the stress response (Figure S1; Supplemental Table 3). During Phase I, upregulated DETs were enriched for terms related to general stress and abiotic stimulus response, water and osmotic stress, translation and ribosome biogenesis, and RNA processing and splicing. Downregulated DETs in Phase I were strongly enriched for photosynthesis, particularly light harvesting and light reactions, consistent with a rapid suppression of photosynthetic activity during turgor loss. ABA-related terms appeared among both Phase I and Phase II upregulated DETs, with the strongest enrichment in Phase II, consistent with the delayed onset of ABA accumulation (Figure 1C). By Phase III, upregulated DETs were enriched for RNA splicing via the spliceosome, translational initiation, and continued water and salt stress responses, suggesting sustained investment in RNA processing and translational reprogramming during stabilisation. The enrichment of RNA processing and splicing terms across multiple phases implies that biosynthesis of the splicing machinery itself is responsive to turgor stress.

### Differential expression revealed only by RNA isoform analysis

We define a Differentially Expressed Gene (DEG) as any gene producing one or more DETs, or any gene for which the total gene expression, the Aggregated Transcript Abundance (ATA), changes significantly. Where a DEG is identified by only one of these criteria, it is referred to accordingly as a DEG_DET_ or DEG_ATA_. We found 13,035 DEG_ATA_ genes (Supplemental Table 4) and 10,167 DEG_DET_ genes (Supplemental Table 2) while 8,909 of these DEGs were identified by both criteria (Figure 2D).

Many DEG_ATA_ genes (4,126) were not identified by DET analysis because individual transcripts did not change significantly but showed similar trends. Conversely, 1,258 DEGs (Supplemental Figure 2) were identified only by analysis of RNA isoforms because changes in a single DET were masked by aggregation with other RNA isoforms from the same gene.

Most intriguingly, 484 genes expressed two DETs with opposing trajectories, one increasing and the other decreasing, such that the ATA did not change. We illustrate this category with four example genes, *P5CS1*, *OSCA1.1*, *KASI*, and *CER3*, each of which has previously been implicated in responses to water deficit (Supplemental Table 1). These genes show minimal net change in expression when transcript abundance is aggregated yet produce RNA isoforms with clearly divergent responses (Figure 2E). Remarkably, such opposing trajectories can be detected for some genes even when they produce large numbers of transcripts. This is illustrated by *P5CS1* for which we detected 29 transcripts (Figure 2E). These results suggest that for hundreds of genes, the response to turgor stress operates through changes in the relative abundance of RNA isoforms rather than overall level of gene expression.

### Turgor stress triggers reversible dominant isoform switching

We next systematically identified all genes with at least two DETs showing opposing trajectories and found 1,704 such genes (Supplemental Figure 3, Supplemental Table 5). In most cases the stress-induced RNA isoform in each opposing pair reached peak expression during Phase II or III (Figure 3A). Isoforms peaking during recovery of turgor (Phase II) might facilitate the restoration of water potential while those peaking later in Phase III might be involved in longer-term responses or stress memory.

**Figure 3.**
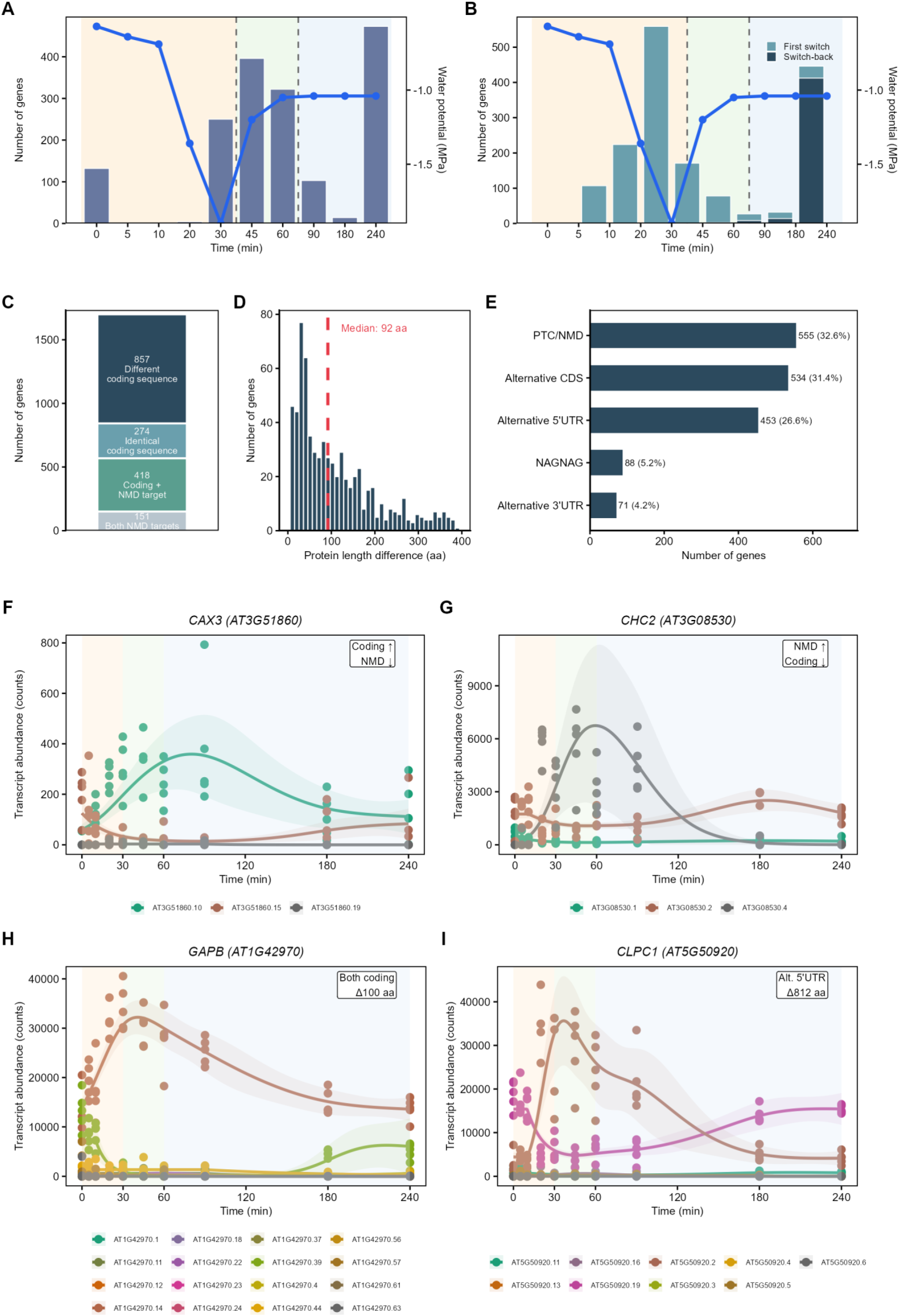
Turgor stress triggers rapid and reversible RNA isoform switching. (A) Timing of peak expression for the stress-induced isoform in each opposing pair, plotted by timepoint. Water potential overlaid (blue line, right axis). Dashed lines indicate phase boundaries. (B) Number of forward switch events (light blue bars) and switch-back events (dark blue bars) at each timepoint. Water potential overlaid as in (A). (C) Predicted coding potential of opposing RNA isoform pairs from 1,704 genes. Categories: both RNA isoforms with different protein-coding sequence (857, dark blue), identical protein-coding sequence (274, light blue), one protein-coding and one predicted NMD-target (418, green), or both predicted NMD targets (155, grey). (D) Distribution of predicted differences in number of amino acids for the 857 genes producing different length proteins. Dashed red line indicates the median difference of 92 amino acids. (E) Classification of structural differences between opposing RNA isoform pairs by mechanism. PTC/NMD: premature termination codon leading to predicted nonsense-mediated decay; Alternative CDS: alternative coding sequence; Alternative 5′ UTR: alternative 5′ untranslated region; NAGNAG: alternative splicing at tandem NAG acceptor sites (Zhang et al., 2022); Alternative 3′ UTR: alternative 3′ untranslated region (F–I) Four examples illustrating distinct functional consequences of isoform switching. (F) *CAX3* (AT3G51860): switch from an NMD-targeted to a protein-coding isoform during stress. (G) *CHC2* (AT3G08530): the reverse pattern, switching from a coding to an NMD-targeted isoform. (H) *GAPB* (AT1G42970): switch between two coding isoforms differing by 100 amino acids. (I) *CLPC1* (AT5G50920): switch between isoforms with alternative 5′ UTRs, resulting in an 812 amino acid difference in predicted protein length. *cpam*-fitted trajectories are shown as described in Figure 1.

For 1,176 of these genes, the response was sufficiently great that the most abundant (dominant) RNA isoform at the beginning declined to become the minor isoform, while an alternative transcript increased to become the dominant isoform. We refer to these as RNA isoform switching events and they occurred predominantly in Phase I, coinciding with the loss in turgor (Figure 3B). This switching was reversed (switch-back events) for many genes and occurred overwhelmingly in Phase III (Figure 3B). This temporal pattern shows that RNA isoform switching is rapid and reversible.

The very rapid switching events reveal that many of the stress-responsive RNA isoforms have very short half-lives, in the order of minutes. Thus, rapid RNA turnover appears to be a feature of the stress response. It is possible that in many cases, one RNA isoform is the precursor of another and is derived by rapid post-transcriptional RNA processing. However, we assume that in many cases, RNA isoforms are produced by changes in transcription.

### Consequences of changes in proportions of RNA isoforms

Of the 1,704 genes with opposing transcript pairs, 857 produced transcripts predicted to encode two different proteins, while only 274 are predicted to encode identical proteins (Figure 3C). A further 418 transcript pairs consisted of one protein-coding transcript and one predicted to be targeted for Nonsense-Mediated Decay (NMD) (Zhang et al., 2022), while for 155 genes both transcripts are predicted to be targets for NMD (Figure 3C). Thus, for the majority of these genes (1,275, 75%), a change in the ratio of RNA isoforms has direct consequences for protein output. In cases where RNA isoforms encoded different proteins, the differences in protein structure were substantial, with the median difference in protein length being 92 amino acids (Figure 3D).

The structural differences between opposing RNA isoforms arises through diverse mechanisms typically involving alternative RNA splicing, alternative transcription start sites or other RNA modifications (Figure 3E). The most common outcome is a frame-shift introducing a premature termination codon (nonsense mutation) leading to predicted NMD (555 genes, 32.6%). For example, the *CAX3* gene shifts from an NMD target to a protein-coding transcript during stress, while *CHC2* shows the reverse pattern (Figure 3F–G). The second category results in alternative coding sequences (534, 31.4%), as illustrated by *GAPB,* which switches between two coding isoforms differing by 100 amino acids (Figure 3H). The third category is characterised by alternative 5’ UTRs (453, 26.6%). One example is *CLPC1* (Figure 3I) which also introduces a new start codon resulting in an 812 amino acid difference in the predicted protein. Smaller proportions were accounted for by NAGNAG splicing (Zhang et al., 2022) and alternative 3’UTRs (Figure 3E).

### Changes in protein domains resulting from RNA isoform switching

Of the 857 genes with opposing trajectory transcripts encoding different proteins, 309 show differences in annotated functional domains of which 216 show a stress induced switch toward a more complete or functionally active isoform (Supplemental Figure 4, Supplemental Table 6). Four examples of genes potentially involved in responses to turgor stress (*OSCA1.1, CDPK2, CCX4, PNET5;* Supplemental Table 1) illustrate this point (Figure 4A). The *OSCA1.1* gene rapidly switches from the RNA isoform containing an inhibitory uORF to one lacking this uORF, then switches back in Phase III. The gene for the stress-responsive calcium-dependent protein kinase CDPK2 switches toward the RNA isoform encoding an intact kinase active site. The gene encoding membrane-located cation/calcium exchanger CCX4 shifts toward the RNA isoform encoding dual ion-binding domains. The gene encoding PNET5 switches toward the RNA isoform encoding a full-length protein including its nuclear envelope-localisation signal (Figure 4A).

**Figure 4.**
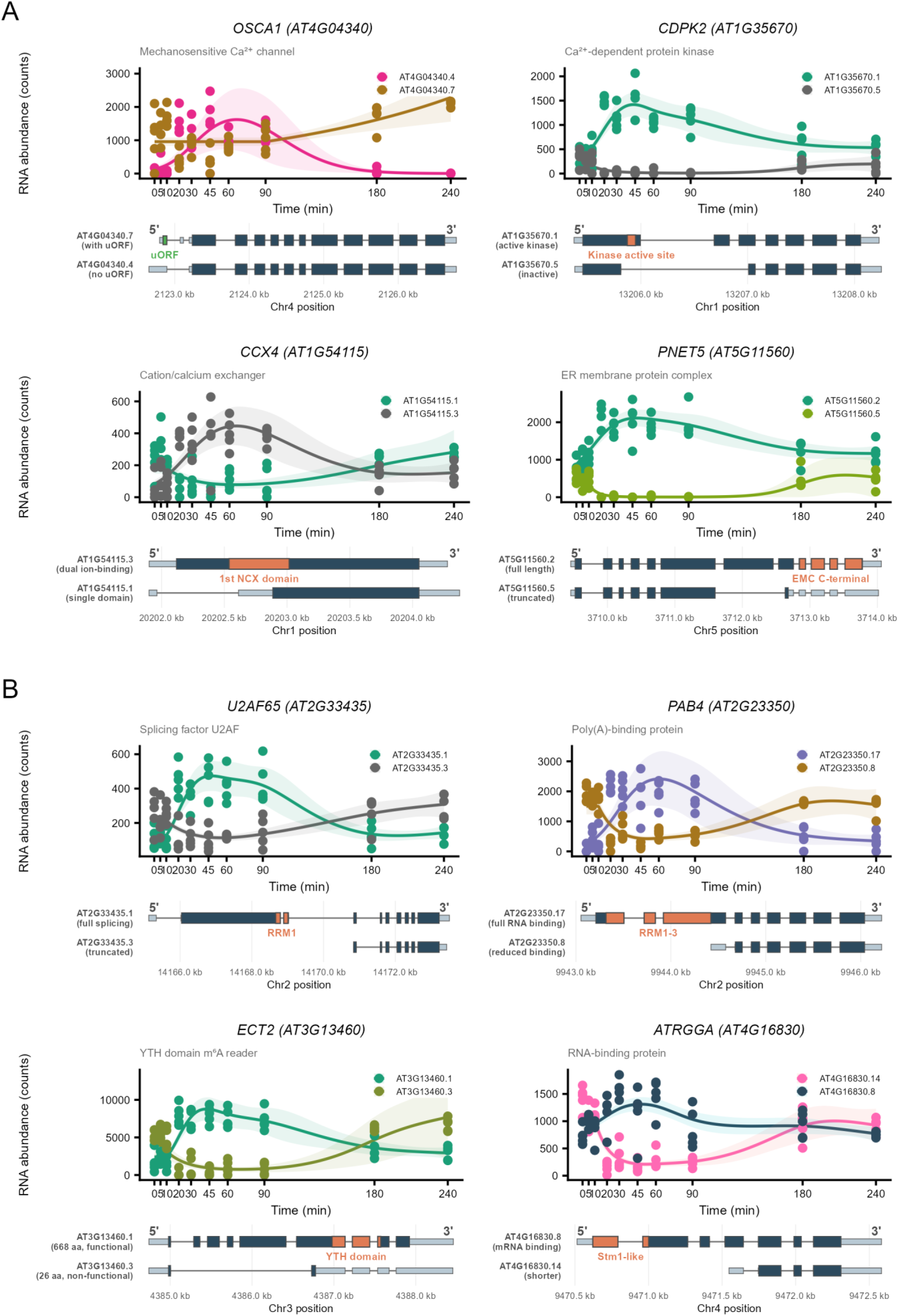
Isoform switching changes protein function in turgor response and RNA processing genes. (A) Changes in expression and predicted coding sequences in RNA isoforms of four turgor response and stress signalling genes. *OSCA1.1* (AT4G04340): mechanosensitive ion channel, isoforms differing by presence of upstream ORF. *CDPK2* (AT1G35670): protein kinase, isoforms differing in kinase active site. *CCX4* (AT1G54115): cation exchanger, isoforms with one or two ion-binding domains. *PNET5* (AT5G11560): nuclear envelope protein, isoforms differing in C-terminal domain including nuclear-envelope signal. The upper plot displays *cpam*-fitted trajectories for only the two transcripts with opposing responses (plotted as described in Figure 1; all other isoforms from each gene are omitted for clarity), and the lower diagram shows the corresponding transcript structures and predicted protein domain differences. In gene structure diagrams, light blue denotes untranslated regions, dark blue denotes coding regions, grey lines represent introns, and orange boxes highlight the variable protein domain distinguishing the two isoforms. Transcript identifiers are indicated alongside each structure (B) Changes in expression and predicted coding sequences in RNA isoforms of four RNA processing and splicing genes. *U2AF65* (AT2G33435): splicing factor, isoforms differing in RRM1 domain. *PAB4* (AT2G23350): poly(A)-binding protein, isoforms with full or reduced RNA-binding domains (RRM1–3). *ECT2* (AT3G13460): m⁶A reader, isoforms of 668 and 26 amino acids. *ATRGGA* (AT4G16830): RNA-binding protein, isoforms differing in Stm1-like domain. Plotted as in (A).

A further set of four examples illustrate the same principle with respect to genes which encode proteins involved in RNA processing splicing, polyadenylation, or mRNA stability (*U2AF65*, *PAB4*, *ECT2,* and *ATRGGA;* Supplemental Table 1). Each example shows a stress-induced switch toward the RNA isoform encoding complete or intact RNA-binding domains (Figure 4B). This observation raises the possibility that changes in the RNA processing machinery which lead to widespread RNA isoform switching, are themselves the result of RNA isoform switching. Consistent with this model, transcripts encoding RNA processing and splicing factors showed significantly earlier changepoints (median 5 min) than those of water and osmotic stress response genes (median 10 min; Wilcoxon p = 5.2 × 10⁻⁹).

### A model for the role of stress-responsive RNA isoforms

At least half the genes in *A. thaliana,* and presumably in other plants, produce more than one RNA isoform. Such RNA isoforms differ in structure and function and so provide the genome with an expanded level of complexity and functional capability. In response to abiotic stresses such as loss of turgor, very rapid post-transcriptional RNA processing mechanisms are brought into play to change the RNA isoform population including RNAs that encode components of the RNA processing apparatus itself. These changes could potentially in turn facilitate further changes in the RNA isoform population as part of the response to the imposed stress. Changes to the RNA isoform population will also come about through changes in transcription. Once the stress is relieved or the plant has adjusted, the RNA isoform population can apparently revert to its previous state or adopt a new, acclimated state.

The extent to which different stresses, such as drought, cold and heat, induce similar changes in the transcriptome are not clear. Analysis of the transcriptome in response to cold stress identified widespread changes in RNA splicing and RNA isoform switching affecting thousands of genes within hours of cold exposure (Calixto et al., 2018; Liu et al., 2022). Several genes switching RNA isoforms in response to cold also switch in response to turgor stress. However, the extent to which different stresses elicit similar changes in the RNA isoform profiles remains to be determined. Similarly, we do not know if different plant species or different ecotypes of the same species employ similar RNA isoforms, or whether plant diversity is in part determined by RNA isoform diversity. These areas of research should be a priority for future studies, to fully understand how plants respond to different environmental stresses. The approaches and extensive datasets presented here provide an important reference point for such studies.

## Materials and methods

### Plant material and growth conditions

*Arabidopsis thaliana* Col-0 seeds were surface-sterilised, stratified at 4°C for 48 h, and sown onto half-strength Murashige and Skoog medium with 0.8% agar, without sucrose (∼100 seeds per plate, spaced to prevent leaf contact with agar or neighbours). Seedlings were grown at 21°C under a 16 h photoperiod (250 µmol m⁻² s⁻¹) for 12 days to the two-true-leaf stage.

### Turgor stress treatment

Plate lids were removed at time zero, exposing seedlings to ambient chamber conditions. Temperature and relative humidity were recorded throughout using a data logger and vapour pressure deficit (VPD) was calculated from these measurements. Chamber temperature was maintained at 21°C. Plates remained stationary in the growth chamber to avoid mechanical stimulation. Seedlings were sampled at 0, 5, 10, 20, 30, 45, 60, 90, 180, and 240 min after lid removal. Only seedlings with no leaf contact with neighbouring plants or the agar surface were sampled. Roots were removed and shoot tissue was immediately frozen in liquid nitrogen. Five biological replicates per timepoint were derived from multiple plates.

### Water potential and ABA measurements

Leaf water potential was measured at each timepoint using thermocouple psychrometers (PSY1, ICT International, Armidale, NSW) as described previously (Ray et al., 2025) with three measurements per timepoint. ABA was quantified from shoot tissue at nine timepoints (0–180 min, three biological replicates) by LC-MS/MS with ²H₆-ABA as an internal standard, following (McAdam, 2015).

### RNA extraction, sequencing and transcript quantification

Total RNA was extracted using the ISOLATE II RNA Mini Kit (Bioline, Ohio). Poly(A)-enriched mRNA libraries were prepared and sequenced by Novogene (Singapore) on the Illumina NovaSeq platform (paired-end 150 bp, ∼6 Gb per sample; 50 libraries total). Transcript abundance was quantified using kallisto (Bray et al., 2016) with 100 bootstrap replicates against AtRTD3 (Zhang et al., 2022). Gene-level abundances were obtained by aggregation using tximport (Soneson et al., 2016).

### Differential expression and changepoint analysis

Temporal differential expression was analysed using the *cpam* R package (Yates et al., 2026). Low-abundance transcripts were filtered using default *cpam* criteria (≥5 reads in ≥60% of samples at one or more timepoints), retaining 95,104 of 154,383 detected transcripts. Quantification uncertainty from kallisto bootstraps was incorporated via overdispersion scaling. Transcripts were classified as differentially expressed at FDR < 0.05, and changepoints were estimated using generalised cross-validation with the one-standard-error rule (Yates et al., 2026)

### Isoform switching analysis

Pairwise Pearson correlations were calculated between all transcript pairs within each multi-isoform gene using mean counts across the ten timepoints. Genes with at least one pair showing r < −0.5 were retained as candidates with opposing trajectories. The dominant isoform at each timepoint was defined as the transcript contributing the highest proportion of total gene expression. A switch was recorded when dominant isoform identity changed between consecutive timepoints with a proportion difference exceeding 10%.

### Functional annotation and GO enrichment

Coding and NMD status were obtained from AtRTD3 annotations (Zhang et al., 2022). Functional domain analysis used InterProScan 5.59-91.0, querying Pfam, SMART, PROSITE and SUPERFAMILY databases. GO enrichment was performed using ShinyGO v0.85 (Ge et al., 2020) with FDR < 0.05, testing DETs separated by phase and direction against all retained transcripts as background. To compare response timing between functional categories, RNA processing/splicing genes and water/osmotic stress genes were defined by GO terms (listed in Supplemental Table 7) and changepoint distributions compared using Wilcoxon rank-sum tests.

## Data Availability

Raw RNA-seq data have been deposited at the NCBI Sequence Read Archive under BioProject accession PRJNA1436459. The *cpam* R package used for changepoint and differential expression analysis is available at https://github.com/l-a-yates/cpam (Yates et al., 2026).

## Author Contributions

JLH and SMS conceived and designed the research. JLH performed experiments. JLH LAY and JBB analysed data. JLH, LAY, JBB and SMS interpreted the data. JLH and SMS wrote the manuscript. JLH, LAY, JBB and SMS edited the manuscript.

## List of Supplemental Figures and Tables

**Supplemental Figure S1. Gene Ontology enrichment analysis of DETs by phase and direction**

**Supplemental Figure S2. Genes identified as differentially expressed through transcript analysis only**

**Supplemental Figure S3. *cpam*-fitted trajectories for all opposing RNA isoform pairs**

**Supplemental Figure S4. *cpam*-fitted trajectories for opposing isoform pairs with predicted changes in protein domain architecture.**

**Supplemental Table S1. Genes referenced in text**

**Supplemental Table S2. Differentially expressed transcripts (DETs)**

**Supplemental Table S3. Gene Ontology enrichment results**

**Supplemental Table S4. Differentially expressed genes identified by aggregated transcript abundance (DEG_ATA_)**

**Supplemental Table S5. Genes with opposing isoform trajectories and switching events**

**Supplemental Table S6. Functional domain differences between opposing isoform pairs**

**Supplemental Table S7. GO terms used for temporal change point comparison**

